# Brain-State-Resolved Consistency of Corticospinal Responses with EEG–TMS

**DOI:** 10.64898/2026.05.25.727597

**Authors:** Timo van Hattem, Juliana R. Hougland, Oskari Ahola, Stefan M. Goetz, Dania Humaidan, Andreas Jooß, Ulf Ziemann

## Abstract

**Background:** Transcranial magnetic stimulation (TMS) over the primary motor cortex (M1) elicits motor-evoked potentials (MEPs), a neurophysiological marker of corticospinal excitability. Ongoing brain activity at the time of stimulation, such as the phase and power of the sensorimotor mu rhythm (8–13 Hz), has a significant impact on MEP amplitudes. However, it remains unclear whether these endogenous excitability states also influence the *consistency* of MEP amplitudes across repeated trials.

**Objectives:** We investigated whether instantaneous mu dynamics modulate not only the magnitude but also the consistency of corticospinal responses to TMS.

**Methods:** Twenty-nine healthy participants received 1200 single TMS pulses over the left M1 during simultaneous EEG recording. Trials were stratified based on pre-stimulus mu power, phase, and interhemispheric M1–M1 functional connectivity. Brain-state-resolved MEP variability was quantified using the coefficient of variation (CV) within subsets of trials defined by similar pre-stimulus mu dynamics.

**Results:** Trial subsets characterized by high mu power or high M1–M1 functional connectivity were associated with reduced MEP variability, indicating more consistent corticospinal output. In contrast, the mu phase did not significantly influence response consistency. Brain-state-resolved MEP variability showed greater stability across sessions compared to MEP variability estimated from random trial subsampling.

**Conclusions:** Pre-stimulus mu dynamics shape not only magnitude but also consistency of corticospinal responses to TMS. We show that corticospinal response consistency reflects a structured, brain-state-dependent property of the sensorimotor network. These findings contribute to our mechanistic understanding of brain-state-dependent neuromodulation and may be leveraged to reduce variability and improve efficacy to TMS.

**Highlights:**

- Ongoing sensorimotor mu dynamics shape both magnitude and consistency of MEPs.
- Trial subsets characterized by high mu power were associated with reduced MEP variability.
- Mu phase modulated MEP amplitude but did not influence MEP consistency.
- Brain-state-resolved estimates of MEP variability were more reliable across sessions.
- Future TMS protocols may reduce effect variability by targeting stable excitability states.

## 1. Introduction

Transcranial magnetic stimulation (TMS) over the primary motor cortex (M1) elicits motor-evoked potentials (MEPs) in the contralateral target muscle, providing a readout of corticospinal excitability [1, 2]. A central challenge in TMS research is the substantial variability in evoked responses, which compromises the reliability and clinical utility of excitability estimates [3, 4, 5]. Trial-to-trial variability in MEPs reflects a combination of methodological (extrinsic) and physiological (intrinsic) sources [6, 7, 8, 9, 10]. Even when methodological parameters are tightly controlled, transient fluctuations in electrophysiological brain state continue to shape the response to TMS [11, 12]. Specifically, pre-stimulus dynamics of the sensorimotor mu rhythm (8–13 Hz), such as the phase and power, modulate corticospinal output [12, 13, 14, 15, 16, 17, 18, 19, 20].

The majority of brain-state-dependent TMS literature focuses on differences in mean MEP amplitude *across* brain states at group level. Trials are typically averaged within state-defined bins to maximize signal-to-noise. The remaining MEP variability after accounting for brain state (i.e., dispersion of MEP amplitudes around the mean within a state-defined bin) is implicitly treated as random, uncontrollable physiological noise. Yet, dispersion of MEP amplitudes *within* a given brain state (henceforth referred to as brain-state-resolved MEP variability or consistency) may itself be an informative feature of the excitability state of the sensorimotor system. To date, only a very limited number of studies have examined response consistency within specific brain states [18], which highlights the need for a more systematic investigation of brain-state-resolved MEP variability.

MEP variability can be modulated by experimental manipulations that increase corticospinal excitability, such as higher stimulation intensities or voluntary muscle contraction, depending on the operating point on the corticospinal input-output (I/O) curve [9, 10, 21, 22]. Whether endogenous high-excitability states, such as the trough and early rising phase of the mu oscillation and high mu power, exert stabilizing effects on corticospinal output similar to externally induced excitability shifts remains unknown.

In the present study, we investigate whether instantaneous pre-stimulus mu dynamics shape not only the magnitude but also the consistency of MEPs. Critically, opposed to assessing variability in terms of mean MEP amplitude between brain states (i.e., between-state MEP variability), we characterize the dispersion of responses once brain state is accounted for (i.e., within-state MEP variability). We quantify brain-state-resolved MEP variability as the dispersion of MEP amplitudes within subsets of trials defined by similar pre-stimulus EEG dynamics, including phase and power, as well as interhemispheric M1–M1 functional connectivity. Identifying brain states in which MEPs are both large in amplitude and more consistent across repeated trials may improve the sensitivity and reliability of corticospinal excitability estimates.

## 2. Methods

### 2.1 Participants

The dataset is comprised of 29 healthy, right-handed participants (18 female; mean ± SD age = 27.4 ± 5.1 years) who underwent a single session of EEG-TMS over the left M1. None of the participants reported a history of neurological or psychiatric disorders or had contraindications to TMS or magnetic resonance imaging (MRI). All participants provided written informed consent prior to participation. The study was approved by the local ethics committee of the University of Tübingen (810/2021BO2) and conducted in accordance with the Declaration of Helsinki.

### 2.2 Experimental protocol

Participants received 1200 single TMS pulses over the hand motor hotspot of the left M1 at 110% of the resting motor threshold (RMT). The motor hotspot was defined as the location that resulted in the most consistent and strongest MEPs in the right abductor pollicis brevis (APB). RMT was determined as the minimum percentage of maximum stimulator output resulting in MEPs with a peak-to-peak amplitude greater than 50 µV in 50% of the trials [23]. During the experiment, participants were seated in a comfortable chair and focused on a fixation cross on a monitor. The session was divided into four blocks of 300 pulses, each block lasting approximately 15 minutes.

### 2.3 Transcranial magnetic stimulation

TMS pulses with a biphasic current waveform were delivered using a figure-of-eight Cool-B65 coil and MagPro R30 stimulator (MagVenture A/S, Farum, Denmark). The randomized inter-trial interval was 2.5–3.5 s. The coil was positioned over the individual hand motor hotspot at approximately 45° relative to the longitudinal fissure. Individual T1-weighted MRI was used for neuronavigation to monitor coil control throughout the session (Localite GmbH, Germany). Individualized noise masking was applied during stimulation blocks [24].

### 2.4 EEG/EMG acquisition

EMG was recorded from the right APB and right first dorsal interosseous (FDI) muscles with a bipolar belly-tendon montage. The TMS-compatible EEG cap consisted of 128 channels with Ag/AgCl ring electrodes (EasyCap, Germany). Data were acquired at 5 kHz using TMS-compatible 24-bit amplifiers with the NeurOne system (Bittium Ltd., Finland). Channels were online referenced to FCz with ground at AFz. Impedances of EEG electrodes were kept < 5 kΩ throughout the session.

### 2.5 Data preprocessing

Data were preprocessed using built-in functions and custom scripts in MATLAB v2024b (MathWorks, Natick, MA, USA) using EEGLAB v2024.2 [25] and the TESA toolbox [26, 27].

#### 2.5.1. EEG/EMG preprocessing

EEG data were epoched from –1.25 to –0.025 s relative to the TMS pulse. Noisy channels were iteratively removed using an automated rejection algorithm based on standardized standard deviation (mean ± SD rejected trials = 8.35% ± 5.31%) [28]. Independent component analysis (ICA) served to suppress ocular artifacts [29, 30]. Data were filtered using a 4th-order zero-phase Butterworth bandpass filter (2–90 Hz) and bandstop filter (48–52 Hz), and down-sampled to 1 kHz. Bad trials in the pre-stimulus EEG were removed (mean ± SD rejected trials = 7.93% ± 4.85%) [28]. Bad channels were spherically interpolated, and all channels were re-referenced to the average reference. Pre-stimulus EEG epochs were cropped to –1.045 to –0.045 s to remove filtering edge artifacts and to minimize potential leakage of post-stimulus activity into the pre-stimulus window due to zero-phase filtering [28]. For further details on the automated rejection algorithm and preprocessing, please refer to Ahola et al. [28].

EMG data were epoched from –100 to 100 ms relative to the TMS pulse, baseline corrected, and linearly detrended using data from the pre-stimulus window (–100 to –10 ms). Trials with pre-innervation, defined as a peak-to-peak EMG amplitude exceeding 50 µV in the pre-stimulus window, were excluded (mean ± SD rejected trials = 4.13% ± 4.45%). MEP peak-to-peak amplitude was extracted by calculating the difference between the first two detected peaks between 20 to 40 ms after TMS onset. Trials without detectable post-stimulus peaks and non-responses, defined as MEP peak-to-peak amplitudes < 50 μV, were excluded to mitigate flooring effects (mean ± SD rejected trials = 4.23% ± 6.12%) [31]. For each participant, the muscle with the highest mean MEP amplitude or best signal quality was selected for further analysis (FDI = 21; APB = 8).

As a quality control measure, trials were excluded if coil position deviated by more than 5 mm or coil orientation deviated by more than 10° relative to the initial recorded motor hotspot (mean ± SD rejected trials = 2.39% ± 5.21%).

### 2.6 Pre-stimulus EEG feature extraction and stratification

Pre-stimulus EEG features were extracted using a surface Laplacian spatial filter centered on electrode C3 (adjacent electrodes: FC5, FC1, CP5, CP1; weights: –0.25) [12, 33]. Mu rhythm (8–13 Hz) band power was computed on a single-trial level using FFT-based spectral power estimation. Functional connectivity within the mu frequency band between the left and right M1 (C3–C4) was quantified using the imaginary part of the phase-locking value (iPLV) [20, 32]. Trials were stratified by dividing the distributions of mu power and M1–M1 functional connectivity into octiles (8 bins) and quartiles (4 bins). Octiles were used to capture continuous trends across brain states, whereas quartiles enabled a more categorical and robust differentiation between low and high states [13, 18]. The lowest (Q1) and highest (Q4) quartiles were respectively labeled as ‘Low’ and ‘High’. A control condition was generated using randomly sampled trial subsets matched in size to each state bin to estimate baseline variability in the absence of stratification by pre-stimulus EEG.

Instantaneous mu phase was estimated using the PHASTIMATE algorithm (autoregressive model order = 30; filter edge size removal = 65; filter order = 128) [34, 35]. For phase estimation, pre-stimulus EEG data were bandpass filtered between 8 and 13 Hz using a Hamming-windowed finite impulse response (FIR) filter. Phase values were binned into four equally sized, centered windows (width π/2) from –π to π (0° = positive peak), corresponding to the trough, rising flank, positive peak, and falling flank [34, 35].

### 2.7 Brain-state-resolved MEP variability quantification

Brain-state-resolved MEP variability was quantified for each subject and state bin using the coefficient of variation (CV):

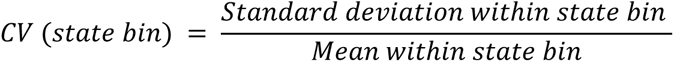

Due to the non-Gaussian, skewed and heteroscedastic nature of MEP distributions (Supplementary Figure 1A-C; Supplementary Table 1) [22, 36, 37], CV was used to express variability relative to the mean within each state and dissociate the effects of these brain states on MEP variability from trivial scaling with the mean. As a robustness check addressing potential mean-variance coupling effects of CV, variability was additionally quantified using the standard deviation (SD) and interquartile range (IQR) of log-transformed MEP amplitudes (see Supplementary Materials) [18].

MEP variability within each state bin was normalized by each participant’s total MEP variability across all trials to account for large inter-individual differences in baseline variability (e.g., anatomy, signal-to-noise ratio, impedance, vigilance) [38]. Normalized CV reflects the proportion of total variability (expressed in percentage) that remains after conditioning on brain state within single individuals.

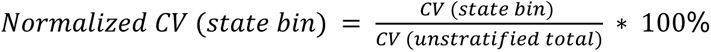

All state bins within each subject contained an equal number of trials to ensure that variability estimates were not confounded by unequal sample sizes [39]. To obtain stable variability estimates, a minimum of 100 usable trials per participant was required following artifact rejection and state stratification. Random subsampling was applied (bootstrapped ten times) to equalize the number of trials across bins within a subject.

### 2.8 Within-session and between-session stability of brain-state-resolved MEP variability

To rule out potential within-session temporal sampling biases (i.e., time-dependent adaptation or vigilance effects), variability differences between mu power bins were examined across the whole session using a sliding-window approach (window size = 300 trials, step size = 50 trials).

In addition, we assessed whether MEP variability represents a stable physiological property across sessions when conditioned on brain states compared to randomly sampled trials. A subset of ten participants (6 female; mean ± SD age 28.5 ± 5.8 years) completed two identical sessions across two separate days. Between-session test-test reliability was quantified using Lin’s concordance correlation coefficient (CCC) [40, 41]. Confidence intervals (CIs) were estimated using non-parametric bootstrapping (n = 1000 with replacement).

### 2.8 Statistical analysis

All statistical analyses were conducted using RStudio v2025.05.1 (R Foundation for Statistical Computing, Vienna, Austria) applying a significance level of α = 0.05. All data are reported as mean ± SEM. For each participant, linear regression was fitted across octiles. Slope coefficients across subjects were tested against zero using two-sided one-sample t-tests against zero. For categorical brain state analyses (i.e., Low vs. High vs. Random power/connectivity, phase), MEP variability was analyzed using a one-factorial repeated measures analysis of variance (rmANOVA) with Greenhouse-Geisser (GG) sphericity correction. Post-hoc pairwise comparisons were performed using paired t-tests with false discovery rate (FDR) correction to control for multiple comparisons.

## 3. Results

### 3.1 High mu power suppresses MEP variability

A progressive reduction in MEP variability was obseved in trial subsets characterized by high mu power (mean ± SEM slope = –1.62 ± 0.41; t(28) = –3.96, p < .001; Figure 1A). In parallel, mean MEP amplitude increased with higher mu power (p < .001; Supplementary Figure 2A). Alternative variability measures confirmed the reduction in MEP variability with increasing mu power (Supplementary Figure 5). Additional control analyses indicated that the observed effects were preferentially associated with the mu frequency range rather than reflecting a general broadband scaling effect (Supplementary Figure 4).

**Figure 1.**
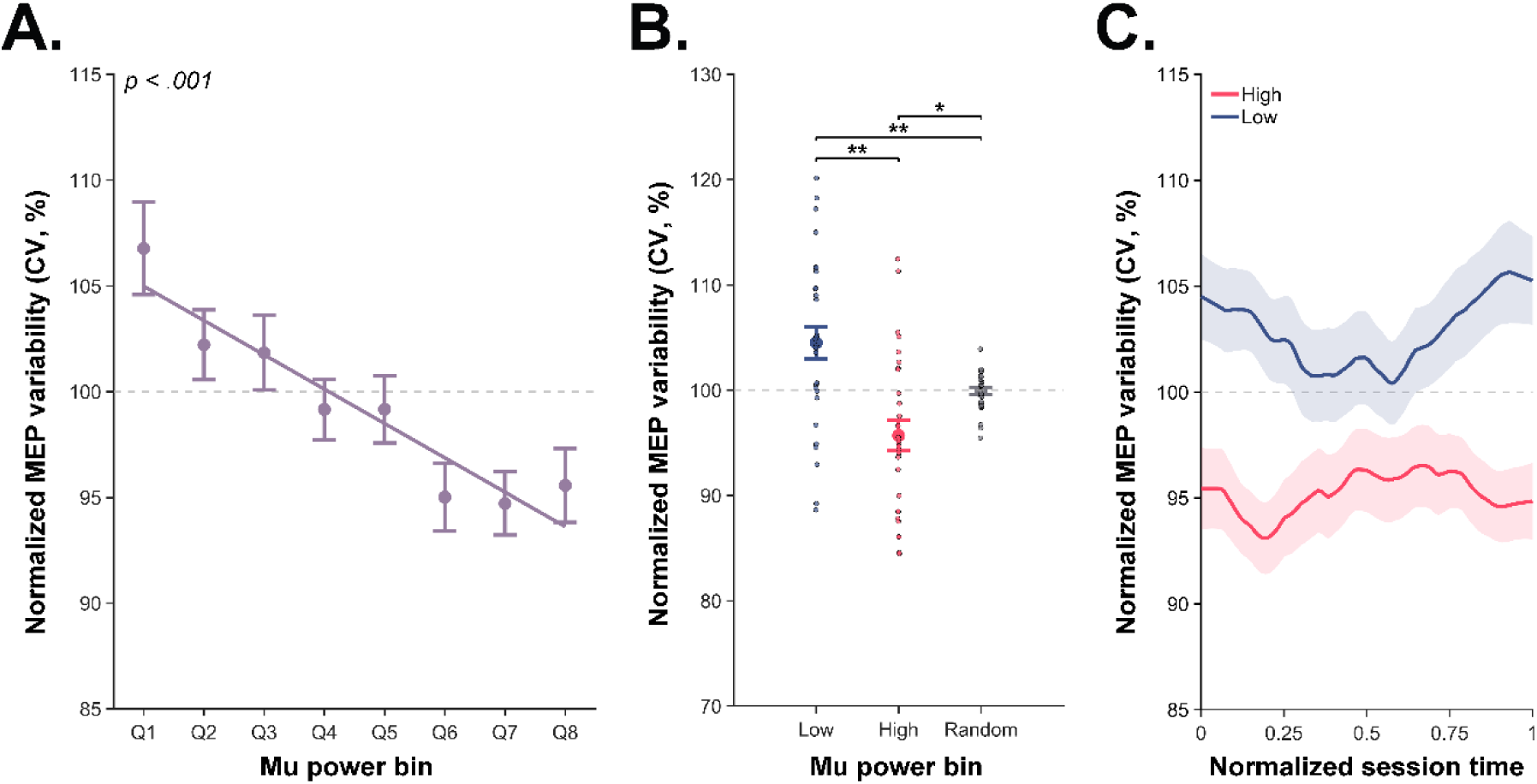
MEP variability resolved by pre-stimulus mu power. **(A)** MEP variability is inversely associated with pre-stimulus mu power across octile percentile bins. **(B)** Comparison of MEP variability in trials subsets with lowest (1st quartile) and highest (4th quartile) mu power. The random condition represents a randomly sampled trial group of equal size. Dots represent data from individual participants. Note the different y-axis range in this subplot compared to the other subplots. **(C)** A sliding window across the experimental session shows consistently lower MEP variability in trials with high mu power compared to trials with low mu power. Session time is normalized from 0 (start of the session) to 1 (end of the session). Y-axes show normalized MEP variability quantified using the coefficient of variation (CV). CV for each state-defined bin is expressed as a percentage of total unstratified CV. The dashed horizontal line at 100% indicates the total unstratified MEP variability (CV across all trials). Error bars and shaded areas represent ± SEM. * p < .05; ** p < .01.

There was a main effect of mu power quartile bins (Low vs. High vs. Random) on MEP variability (F(1.28, 35.83) = 10.09, p = .002; Supplementary Table 1) with post-hoc comparisons showing that MEP variability was significantly lower in the high-power bin compared to low-power bin (t(28) = 3.39, p = .006; Figure 1B). Relative to the random, MEP variability was significantly higher in low-power bin (t(28) = 2.99, p = .009; Figure 1B), and lower in high-power bin (t(28) = −2.69, p = .012; Figure 1B). The observed relationship between mu power and MEP variability was stable throughout the experimental session (Figure 1C).

### 3.2 Mu phase does not modulate MEP variability

There was no significant group-level effect of mu phase on MEP variability (F(2.93, 82.04) = 1.77, p = .160; Figure 2A; Supplementary Table 1). In contrast, mu phase did significantly modulate mean MEP amplitude (p = .002; Supplementary Figure 2B). Control analyses using alternative variability estimates (Supplementary Figure 5) and alternative phase estimation parameters yielded comparable null effects (Supplementary Figure 3).

**Figure 2.**
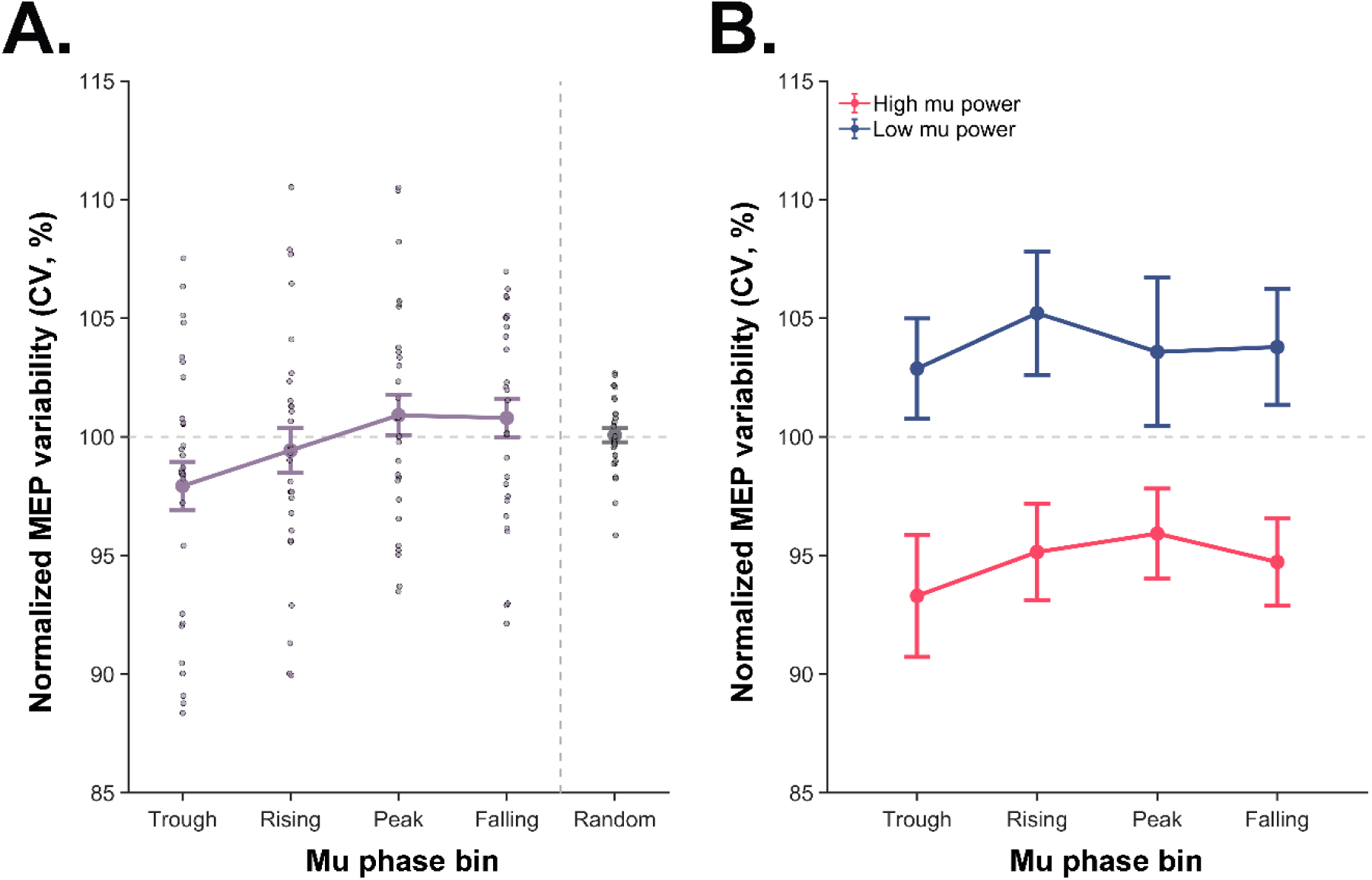
MEP variability resolved by mu phase. **(A)** Mu phase across four phase bins does not significantly modulate MEP variability on group-level. **(B)** Stratifying trials by mu power does not reveal a phase-dependent modulation of MEP variability. Y-axes show normalized MEP variability quantified using the coefficient of variation (CV). CV for each state-defined bin is expressed as a percentage of total unstratified CV. The dashed horizontal line at 100% indicates the total unstratified MEP variability (CV across all trials) against which within-state variability was normalized. Dots represent data from individual participants. Error bars represent ± SEM. * p < .05; ** p < .01; *** p < .001.

To assess whether phase effects on MEP variability emerges within specific mu power states, MEP variability was assessed using joint stratification of mu power bin and phase. There was a significant main effect of mu power on MEP variability (F(1, 28) = 12.42, p = .001; Figure 2B; Supplementary Table 1), but no main effect of phase, (F(3, 84) = 0.37, p = .772; Figure 2B), and no interaction (F(3, 84) = 0.12, p = .951; Figure 2B).

### 3.3 Modulation of MEP variability by interhemispheric M1–M1 functional connectivity

There was a significant reduction in MEP variability associated with trial subsets with higher interhemispheric M1–M1 connectivity strength (mean ± SEM slope = –0.82 ± 0.25, t(28) = -−3.29, p = 0.003; Figure 3A). M1–M1 connectivity did not significantly affect mean MEP amplitude (p = 0.055; Supplementary Figure 2C).

**Figure 3.**
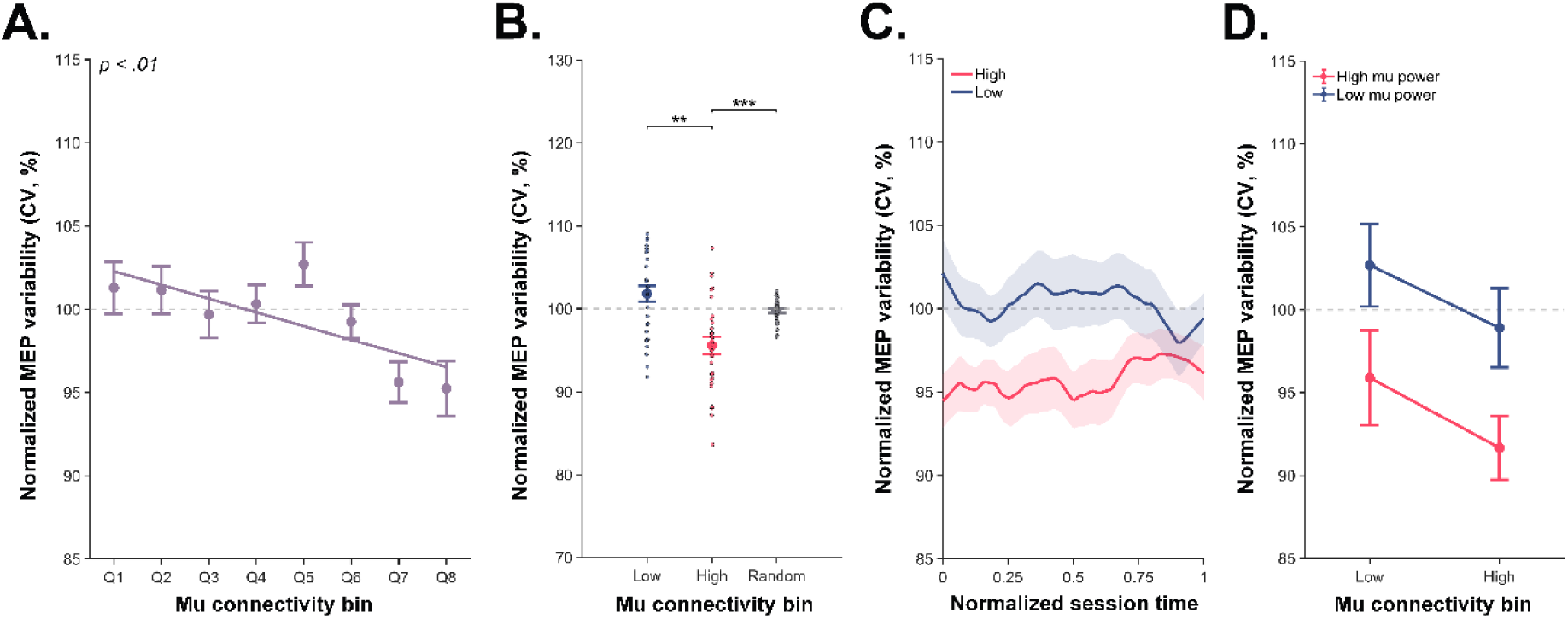
Modulation of MEP variability by interhemispheric M1–M1 functional connectivity. **(A)** Increasing interhemispheric M1–M1 connectivity in the mu frequency band is associated with reduced MEP variability. **(B)** MEP variability differs across connectivity states in comparison to random subsampling. Dots represent data from individual participants. Note the different y-axis range in this subplot compared to all other subplots. **(C)** Within-session temporal stability analysis shows that connectivity-dependent differences in MEP variability remain relatively consistent across the experimental session. **(D)** Joint stratification by mu power and M1–M1 connectivity shows an additive effect on MEP variability with lowest variability during high-power/high-connectivity and highest variability during low-power/low-connectivity. Y-axes show normalized MEP variability quantified using the coefficient of variation (CV). CV for each state-defined bin is expressed as a percentage of total unstratified CV. The dashed horizontal line at 100% indicates the total unstratified MEP variability (CV across all trials) against which within-state variability was normalized. Error bars and shaded areas represent ± SEM. ** p < .01; *** p < .001.

A main effect of mu connectivity bin (High vs. Low vs. Random) on MEP variability was observed (F(1.26, 35.21) = 12.35, p < .001; Supplementary Table 1). Post-hoc comparisons showed that during high M1–M1 connectivity, MEP variability was significantly lower compared to low M1–M1 connectivity (t(28) = 3.67, p = .002; Figure 3B) and lower compared to the random bin (t(28) = –4.35, p = < .001; Figure 3B). The relationship between M1–M1 connectivity and MEP variability was stable throughout the experimental session (Figure 3C).

Combined stratification demonstrated a significant main effect of mu power (F(1,25) = 5.91, p = .023; Supplementary Table 1), but no significant main effect of mu M1–M1 connectivity (F(1,25) = 2.92, p = .10) or interaction (F(1,25) = 0.01, p = .92; Figure 3D). MEP variability was significantly lower in the high-power/high-connectivity condition compared to the low-power/low-connectivity condition (t(48) = 2.96, p = .03; Figure 3D).

### 3.4 Between-session test-retest reliability of MEP variability resolved by mu power

The effect of mu power on MEP variability was reproducible across sessions at both the group level (Figure 4A) and within subjects (Figure 4B). There was a significant main effect of state, F(1.18, 10.59) = 6.22, p = .027; Figure 4A), while no significant main effect of session (F(1, 9) = 0.02, p = .900) or interaction (F(1.84, 16.52) = 0.14, p = .853) was found.

**Figure 4.**
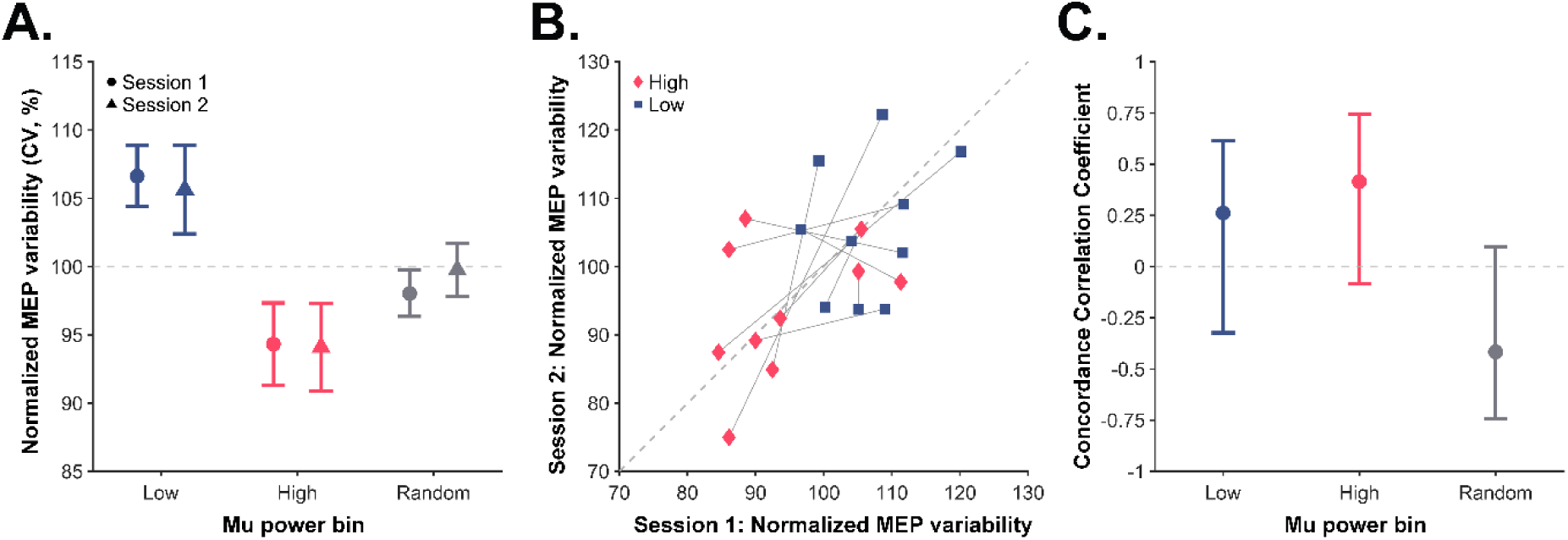
Test-retest of MEP variability resolved by mu power. (A–B) Ten participants had two sessions with the same stimulation protocol across separate days. Effect of mu power on MEP variability was consistent across experimental sessions on a group-level (A) and subject-level (B). Y-axes show normalized MEP variability quantified using the coefficient of variation (CV). CV for each state-defined bin is expressed as a percentage of total unstratified CV. The dashed horizontal line at 100% in (A) indicates the total unstratified MEP variability (CV across all trials) against which within-state variability was normalized. Error bars represent ± SEM. The identity line in (B) indicates identical measures of MEP variability in Session 1 and 2. **(C)** MEP variability shows moderate test-retest reliability (High: 0.42; Low: 0.26) across sessions when trials are resolved by mu power, as quantified by Lin’s concordance correlation coefficient (CCC). In contrast, random trial grouping yields negative CCC estimates, indicating poor reliability under random trial sampling. Error bars represent ± 95% confidence intervals. The 95% confidence interval was defined as the 2.5th and 97.5th percentiles of the bootstrap distribution.

Mu power-resolved MEP variability showed weak-to-moderate test-retest reliability, with positive CCC values observed for both the low-power bin (CCC = 0.26, 95% CI = [-0.32, 0.61]; Figure 4C) and high-power bin (CCC = 0.42, 95% CI = [-0.08, 0.75]; Figure 4C). In contrast, random trial grouping yielded a negative CCC estimate (CCC = –0.42, 95% CI = [-0.74, 0.10]; Figure 4C).

## 4. Discussion

The present study investigated whether ongoing sensorimotor mu rhythm dynamics influence not only the magnitude but also the consistency of MEP amplitudes. We found that trial subsets characterized by high mu power and high interhemispheric M1–M1 functional connectivity were associated with reduced MEP variability. In contrast, the instantaneous phase of the mu rhythm did not significantly modulate response consistency. Together, these findings demonstrate that the consistency of corticospinal output across trials is a structured, state-dependent feature of the sensorimotor system.

### 4.1 Brain-state-dependent modulation of response magnitude versus response consistency

Previous EEG-TMS studies have demonstrated that the instantaneous power and phase of the sensorimotor mu rhythm are linked to trial-to-trial fluctuations in MEP amplitudes [e.g., 12, 14, 15, 16, 17]. The present study replicates these group-level effects (Supplementary Figure 2). Our findings expand prior work by demonstrating that mu rhythm dynamics may also regulate response consistency across trials in addition to response magnitude. The inverse relationship between mu power and interhemispheric M1–M1 functional connectivity with MEP variability suggest that cortical network dynamics contribute to stabilizing corticospinal responses. The absence of a detectable effect of mu phase on MEP variability indicates that phase-dependent modulation influences corticospinal response magnitude rather than consistency.

### 4.2 Response consistency through cortico-motoneuron recruitment gain

Mathematical models have been developed to characterize the sources of MEP variability, describing how trial-to-trial fluctuations emerge from threshold-dependent cortico-motoneuron recruitment interacting with noisy synaptic input currents (i.e., intrinsic variability) [22, 36, 37, 42, 43, 44]. In the present study, stimulation was consistently applied at 110% RMT, corresponding to the point of maximal dispersion around the rising slope of the recruitment curve [22, 36]. This non-saturated operating regime thus maximizes sensitivity to brain-state-resolved variability across trials, which may affect whether some neurons near their threshold respond or not, and minimizes confounding effects of stimulation intensity.

At a fixed stimulation intensity, residual MEP variability is attributed to fluctuations in effective synaptic input and motoneuron pool recruitment [9, 36]. At lower levels of cortico-motoneuron recruitment, corticospinal output is thought to rely more strongly on the probabilistic activation of a relatively small subset of neurons near firing threshold [36]. In this global low-recruitment regime, small fluctuations in membrane excitability or synaptic input could lead to disproportionately large changes in output, resulting in higher variability across trials. In contrast, when a larger pool of neurons is recruited uniformly, the contribution of individual threshold fluctuations may partially average out across the population. Consequently, stochastic fluctuations in effective synaptic input have proportionally smaller influence on the MEP amplitude, leading to a more stable response and reduced variability across repeated trials [36].

### 4.3 Large-scale network states as endogenous modulators of corticospinal recruitment

Mu power reflects a population-level state that indexes the degree of neuronal firing within the sensorimotor network [45]. Thus, TMS interacts with a broader and more uniformly engaged neuronal population during high-power states. Following Capaday’s threshold-based model [36], mu power may act as an endogenous driver that shifts the operating point of the corticospinal system toward a more saturated recruitment regime. As MEP generation becomes less dominated by stochastic threshold effects [36], MEP amplitudes may become larger in size *and* less variable across trials. One possible interpretation is that in this situation organized population activity exerts a stronger influence on corticospinal output and the relative contribution of intrinsic background fluctuations to trial-to-trial variability is reduced. Importantly, intrinsic physiological noise remains present, but its relative impact may be diminished when power is high because a larger proportion of the cortico-motoneurons are consistently driven close to firing threshold. They will more consistently fire an action potential to TMS compared to a low-power state where a larger proportion of cortico-motoneurons is further away from firing threshold and will require additional synaptic input to reach the firing threshold. This additional synaptic input is variable according to the intrinsic background fluctuations in brain state. Thus, the observed effects may refer to a relative signal-to-noise phenomenon rather than absolute suppression of neural noise.

Similarly, stronger interhemispheric functional connectivity may reduce MEP variability by promoting more coordinated neuronal activity across distributed cortical regions. This increased coupling could provide more coherent and synchronized synaptic input to corticospinal neurons, thereby increasing the proportion of consistently recruited neurons and reducing the relative influence of stochastic fluctuations on corticospinal output. In combination, MEP variability decreased linearly across joint mu power and connectivity trial subsets, reaching a minimum when power and connectivity was high. This suggests that large-scale network properties such as power and interhemispheric functional connectivity can jointly stabilize corticospinal output. This pattern matches recent intracranial electrical stimulation work showing that pre-stimulus network dynamics can control the variability of post-stimulus responses [46].

### 4.4 Null effect of mu phases on MEP consistency

Although sensorimotor mu phase significantly modulated mean MEP amplitude, it did not significantly influence MEP consistency across repeated trials. This lack of a detectable effect of the mu phase on MEP variability at the group-level aligns with previous research that reported no significant modulation of MEP variability by only a selected number of phase-power conditions [18].

The absence of effects on MEP consistency may reflect the transient nature of phase-dependent excitability modulation. Mu phase likely introduces only a brief temporal bias in sensorimotor excitability rather than a sustained change in corticospinal recruitment. Thus, although phase may influence mean MEP amplitude within narrow oscillatory windows, it may not sufficiently overcome the relative contribution of ongoing background noise to corticospinal output across repeated trials. Moreover, MEP variability arises from multiple stochastic corticospinal and peripheral sources [22, 37, 42]. These downstream, non-oscillatory fluctuations are likely only weakly phase-locked to cortical rhythms and, therefore, remain largely unaffected by the timing of stimulation within an oscillatory cycle.

Importantly, mu phase may represent a relatively unstable excitability state that fluctuates across time and sessions [47], potentially limiting its ability to reliably constrain corticospinal variability across many trials acquired during a long experimental session. Moreover, the effect sizes of mu phase on corticospinal excitability are generally modest to weak (Supplementary Figure 1B) [48, 49], which indicates that mu phase may not fully capture the full excitability state of the corticospinal system. Consistent with this, previous studies have reported substantial interindividual variability in phase-dependent effects, with many participants showing little or no significant mu phase modulation of corticospinal excitability [47, 48, 50]. Interindividual differences in mu phase sensitivity may also have contributed to the observed modest variability effects (Supplementary Figure 2).

### 4.5 Cross-site generalizability and variability metric sensitivity

To evaluate the generalizability of the findings, we included a validation dataset acquired at Aalto University (Finland) using a similar EEG-TMS protocol over the left M1. Despite minor hardware and recording differences (see Supplementary Materials), findings replicated across sites (Supplementary Figure 6), suggesting that brain-state-dependent modulation of MEP variability reflects a robust physiological effect rather than driven by site-specific methodology.

In the present study, MEP variability was primarily quantified using the coefficient of variation (CV), consistent with prior work [9, 51, 52]. However, CV may be sensitive to distributional shape, outliers, and mean-dependent scaling effects. To address this, we complemented CV with quantification analyses that are less dependent on the mean and correct for distributional skewness, including the standard deviation and interquartile range of log-transformed amplitudes [18]. Power and phase effects on MEP variability were qualitatively robust across all metrics (Supplementary Figure 5A-F).

### 4.6 Implications for brain-state-informed TMS–EEG readouts

Variability in TMS-evoked responses undermines reproducibility, reliability and clinical utility of TMS-based biomarkers and interventions [3, 4, 5, 53, 54]. Our findings suggest that a proportion of physiological variability in corticospinal responses is structured and brain-state-dependent. These results support using real-time EEG-informed TMS to further control and reduce outcome variability. In particular, targeting endogenous “consistent excitability states” associated with reduced variability *and* higher excitability (e.g., high mu power states) may improve the robustness and prognostic value of TMS biomarkers [4, 55]. This may be especially relevant in pre-post intervention paradigms, where fluctuations in baseline corticospinal state can obscure genuine plasticity effects related to a therapeutic TMS intervention.

The reliability of mu power-resolved MEP amplitude variability is demonstrated in our between-session test-retest analyses (Figure 4). The negative concordance observed for randomly selected trials suggests that the MEP variability is dominated by session-specific sources of variability, including differences in vigilance, electrode placement, and background EMG activity. In contrast, conditioning variability estimates on pre-stimulus brain state substantially improved concordance across sessions. This may indicate that conditioning on brain state partially controls the total variability, reducing between-session heterogeneity across days. Importantly, reliability remained weak-to-moderate (with relatively wide confidence intervals overlapping zero), consistent with the presence of residual non-neural and neural sources of variability.

Our findings show that while two brain states may produce similar mean MEP amplitudes, they may yet differ markedly in how consistently they respond across trials. This raises the possibility that optimal brain states for therapeutic TMS interventions are not necessarily those associated with the largest responses, but those that combine higher excitability with greater response consistency. Although speculative at this point, this could potentially provide a stronger net cumulative plasticity induction and more reliable intervention effects across a stimulation session. Future closed-loop brain-state-dependent TMS protocols may benefit from targeting such “consistent high-excitability states”.

Beyond controlling variability, brain-state-resolved response consistency itself may carry clinically relevant biomarker information, as it reflects the stability of corticospinal recruitment and may index corticospinal dysfunction in neurological conditions such as chronic stroke [56, 57, 58].

### 4.7 Limitations

Although the coefficient of variation (CV) accounts for differences in mean MEP amplitude across brain states, it may remain partially influenced by residual mean-variance coupling. While supplementary analyses using alternative variability metrics produced qualitatively similar findings, we cannot fully exclude the possibility that part of the observed brain-state-dependent variability reflects statistical dependence on mean amplitude rather than entirely independent modulation of response consistency. Future studies explicitly modeling mean-variance relationships may help disentangle these effects more precisely.

Finally, estimates of variability may be influenced by preprocessing choices and analysis pipeline decisions. Methodological factors such as filtering parameters, target muscle selection, phase estimation procedures, and approaches to spectral decomposition can affect trial-wise brain state classification and the resulting variability measures [59].

## 5. Conclusions

We quantified how pre-stimulus sensorimotor mu rhythm dynamics shape variability in corticospinal output. MEP variability was primarily modulated by mu power state and interhemispheric M1–M1 functional connectivity, but not strongly by mu phase. These findings show that corticospinal variability contains a reproducible brain-state-dependent component that generalizes across sessions and independent datasets. Together, the results advance our mechanistic understanding of brain-state-dependent neuromodulation and may improve the consistency and efficacy of TMS protocols.

## CRediT authorship contribution statement

**Timo van Hattem:** Conceptualization, Project administration, Methodology, Formal analysis, Visualization, Writing – original draft, Writing – review & editing. **Juliana R. Hougland:** Validation, Formal analysis, Writing – review & editing. **Oskari Ahola:** Investigation, Writing – review & editing. **Stefan M. Goetz:** Writing – review & editing. **Dania Humaidan:** Supervision, Writing – review & editing. **Andreas Jooß:** Supervision, Writing – review & editing**. Ulf Ziemann:** Funding acquisition, Resources, Supervision, Writing – review & editing.

