## Supplementary Materials for "Brain-State-Resolved Consistency of Corticospinal Responses with EEG–TMS"

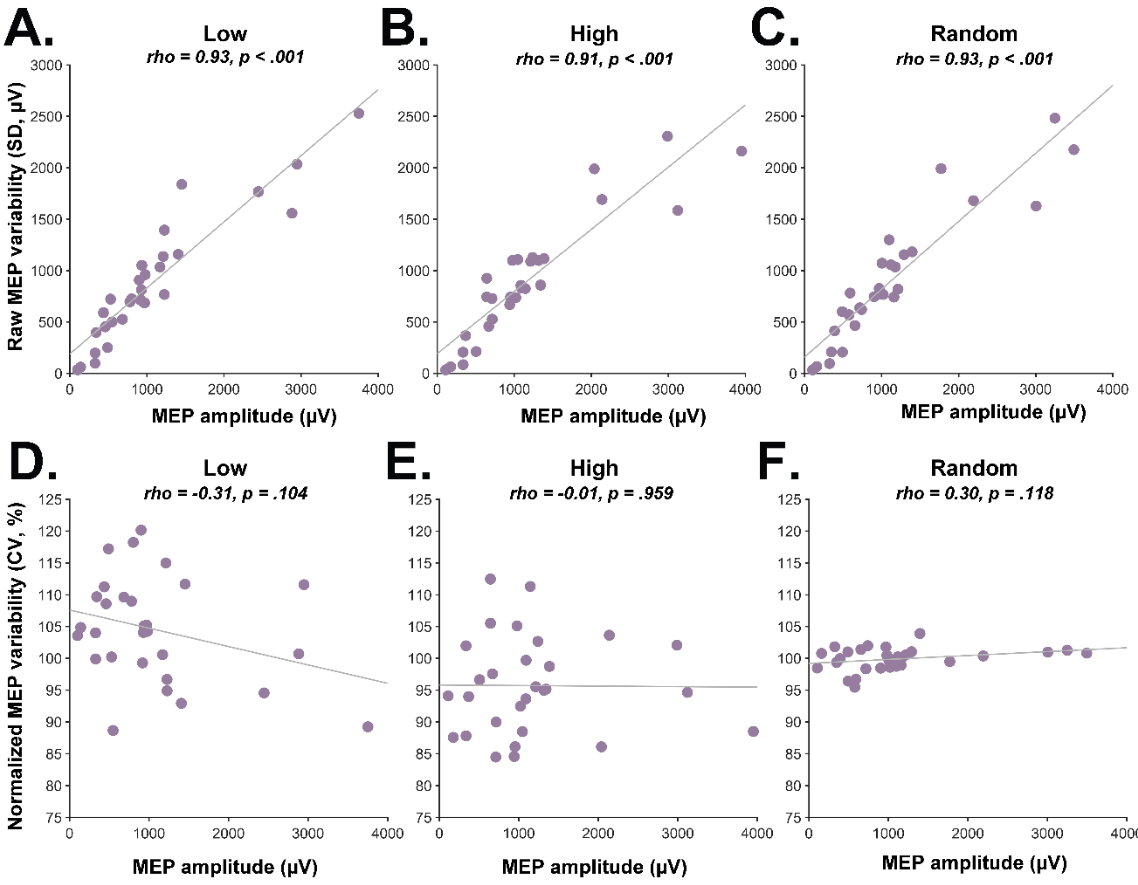


**Supplementary Figure 1. Relationship between MEP amplitude and MEP variability across mu power bins. (A-C)** Correlations across subjects between mean MEP amplitude and absolute MEP variability (raw SD of MEP amplitudes) for the Low, High and Random mu power trial subsets. Significant positive correlations were observed, indicating that subjects with larger mean MEP amplitudes also show greater raw MEP variability, consistent with amplitude-dependent heteroscedasticity. **(D-F)** Correlations across subjects between mean MEP amplitude and normalized MEP variability (CV) for the Low, High, and Random mu power trial subsets. No significant relationships were observed, indicating that the reduction in variability after resolving by mu power was not explained by differences in mean MEP amplitude. Together, these findings show that although raw MEP variability scales with MEP magnitude, the reduction in MEP variability observed after resolving by mu power reflects a genuine stabilization of responses rather than a trivial consequence of differences in mean amplitude.

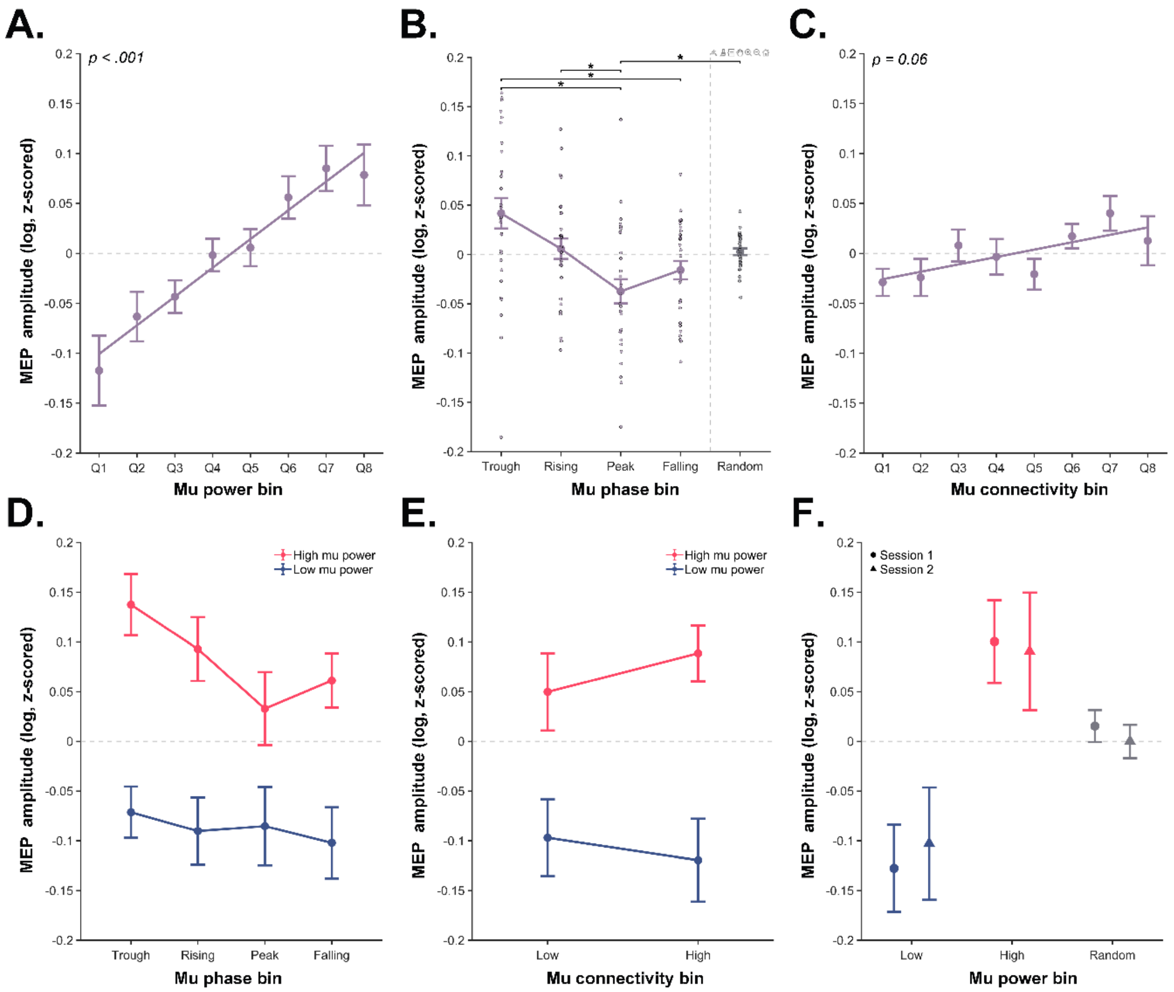


**Supplementary Figure 2. Brain-state-dependent modulation of MEP magnitude.**
**(A)** Mean MEP amplitude was positively associated with mu power across octile percentile bins, which indicates a significant increase in MEP amplitude in trial subsets with higher mu power (mean ± SEM slope = 0.03 ± 0.01; $t(28) = 3.93$, $p < .001$). (B) Mu phase significantly modulated MEP amplitude ($F(2.37, 66.23) = 6.03$, $p = .002$, $\eta^2 = .001$), with the trough associated with the highest MEP amplitudes and the peak with the lowest. At the individual level, ~14% of participants showed a significant mu phase effect on MEP amplitude using circular-to-linear correlation [1], consistent with earlier reports [2, 3]. (C) Connectivity showed a positive association with mean MEP amplitude, but this effect did not reach statistical significance (mean ± SEM slope = 0.007 ± 0.004, $t(28) = 2.01$, $p = .055$). (D) The effect of phase on MEP amplitude in low mu power regimes (1st quartile) and high mu power regimes (4th quartile). (E) The combined effect of mu power and M1–M1 connectivity on mean MEP amplitude. (F) Between-session test-retest of mu power on mean MEP amplitude. Y-axes show z-scored log-transformed peak-to-peak MEP amplitudes. The dashed horizontal line indicates log transformed z-scored MEP amplitude = 0. Error bars indicate ± SEM. $p < .05$.

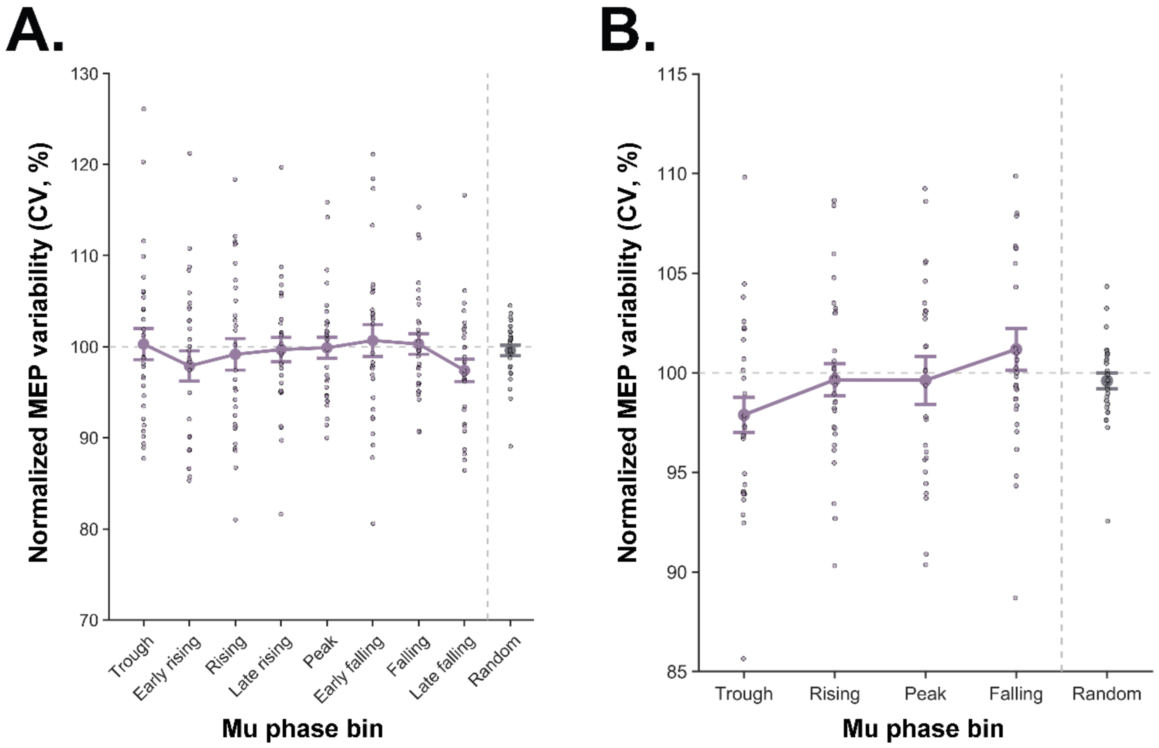


**Supplementary Figure 3. Mu phase-resolved MEP variability using alternative phase estimation approaches. (A)** When using eight mu phase bins [4, 5], no significant main effect of phase on MEP variability was observed ($F(5.71, 159.80) = 0.57$, $p = .748$). (B) When using alternative preprocessing with a narrower phase estimation window and using individual mu peak frequencies [6], no significant main effect on MEP variability was observed ($F(2.67, 74.86) = 1.40$, $p = .253$). The mu-peak frequency was defined as the frequency with the highest power within the mu band (8–13 Hz). Using the PHASTIMATE toolbox, pre-stimulus EEG signal that was epoched [-1.005, -0.005 s] was filtered using an 80th-order FIR bandpass filter (stopband: mu peak ± 5 Hz; passband: mu peak ± 1 Hz). The instantaneous phase at the time of the TMS pulse was extracted (filter order = 128, edge samples removed = 40, autoregressive model order = 20) [5, 6]. Y-axes show z-scored log-transformed peak-to-peak MEP amplitudes. The dashed horizontal line at 100% indicates the total unstratified MEP variability (CV across all trials) against which within-state variability was normalized. Dots represent data from individual participants. Error bars indicate ± SEM.

### Frequency specificity and broadband scaling effect

In this study, power was defined using band-limited mu without explicitly dissociating oscillatory components from broadband spectral activity. Although this approach is similar to prior brain-state-dependent TMS studies [7, 8, 9], band power measures may partly reflect global factors such as vigilance or internal cognitive state, rather than a band-specific neural effect. We performed a sensitivity analysis using mu power normalized by total band power, which yielded comparable main effects (Supplementary Figure 4D). Additionally, separation of periodic mu power from aperiodic mu power showed that variability is constrained by both the periodic and aperiodic mu component (Supplementary Figure 4B). Notably, no comparable effects were observed in other frequency bands, arguing against a general broadband scaling effect (Supplementary Figure 4C-D).

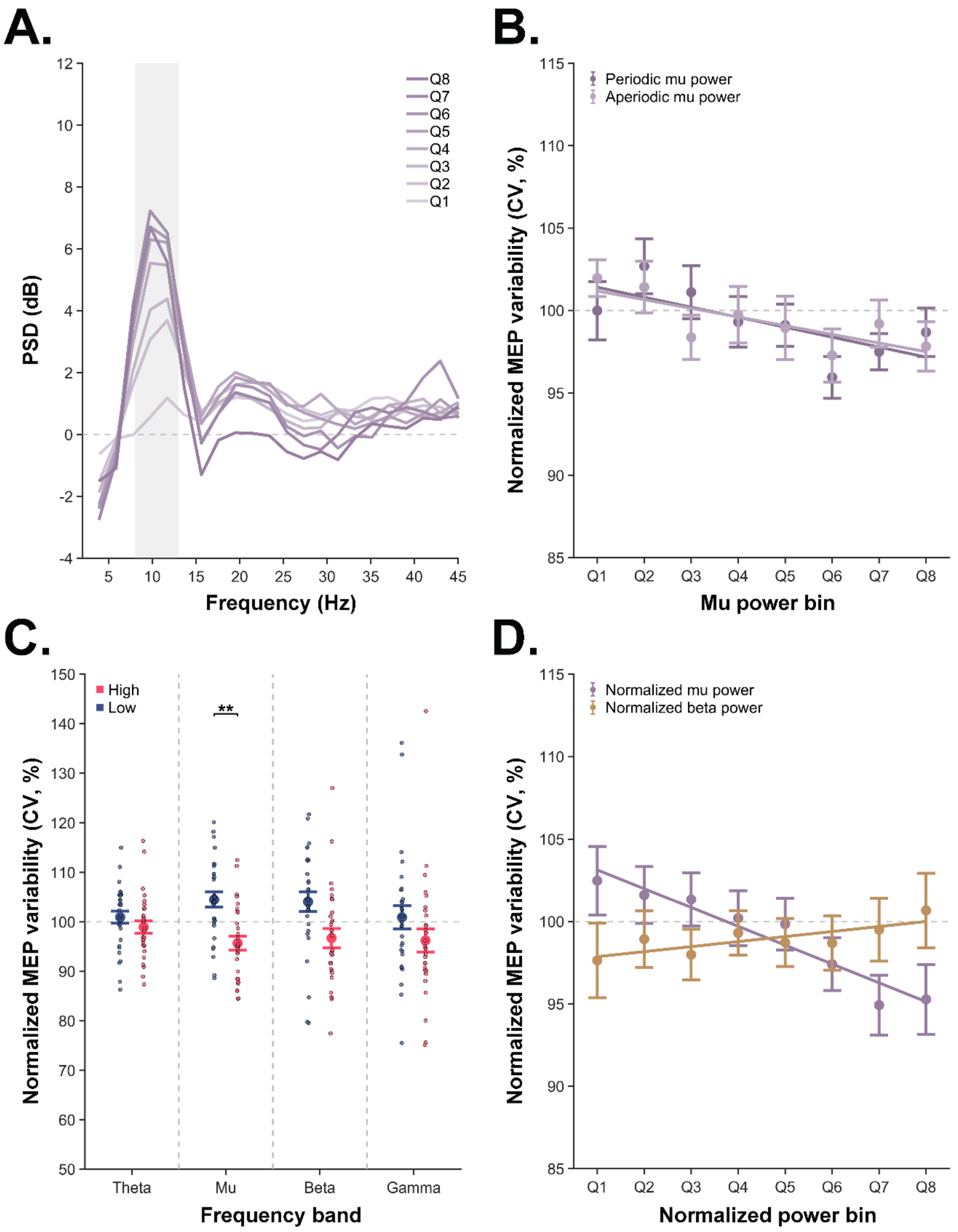


**Supplementary Figure 4. Specificity of mu power effects on MEP variability. (A)** Mean power spectral density plot (1/f-corrected) across the eight mu power bins. (B) Decomposition of mu power into periodic and aperiodic components using the FOOOF algorithm at the single-trial level (peak width limits: (2, 8); maximum number of peaks: 4; minimum peak height: 0.1) [6] revealed that both periodic (mean ± SEM slope = –0.60 ± 0.31; $t(28) = –1.95$, $p = .061$) and aperiodic mu power (mean ± SEM slope = –0.53 ± 0.28; $t(28) = –1.88$, $p = .072$) may contribute to the observed reduction in MEP variability. (C) Extending the model to include other frequency bands (theta [4–8 Hz], beta [13–30 Hz], and gamma [30–80 Hz]) revealed a significant main effect of power state (Low vs. High) ($F(1,28) = 6.02$, $p = .021$), but no main effect of frequency band ($F(2.05, 57.33) = 1.35$, $p = .268$) and no interaction ($F(1.88, 52.55) = 1.33$, $p = .273$). Paired t-tests within each frequency band showed that only mu power survived multiple-comparison correction (theta: $t(28) = –0.91$, $p = .367$; mu: ($t(28) = –3.39$, $p = .008$; beta: $t(28) = -2.05$, $p = .101$; gamma: $t(28) = –1.11$, $p = .366$), suggesting a frequency-specific effect rather than global broadband effects. (D) Normalizing mu band power to total band power to control for global (broadband) fluctuations yielded consistent results for mu power (mean ± SEM slope = –1.14 ± 0.48; $t(28) = –2.37$, $p = .025$), which suggests that the effect is not solely driven by broadband power changes. Any effect of beta power appeared to be driven by broadband power scaling as it did not hold up when normalizing by total band power (mean ± SEM slope = 0.30 ± 0.39; $t(28) = 0.77$, $p = .445$). Y-axes show normalized MEP variability quantified using the coefficient of variation (CV). CV for each state-defined bin is expressed as a percentage of total unstratified CV. The dashed horizontal line at 100% indicates the total unstratified MEP variability (CV across all trials) against which within-state variability was normalized. Dots represent data from individual participants. Error bars represent ± SEM. ** $p < .01$.

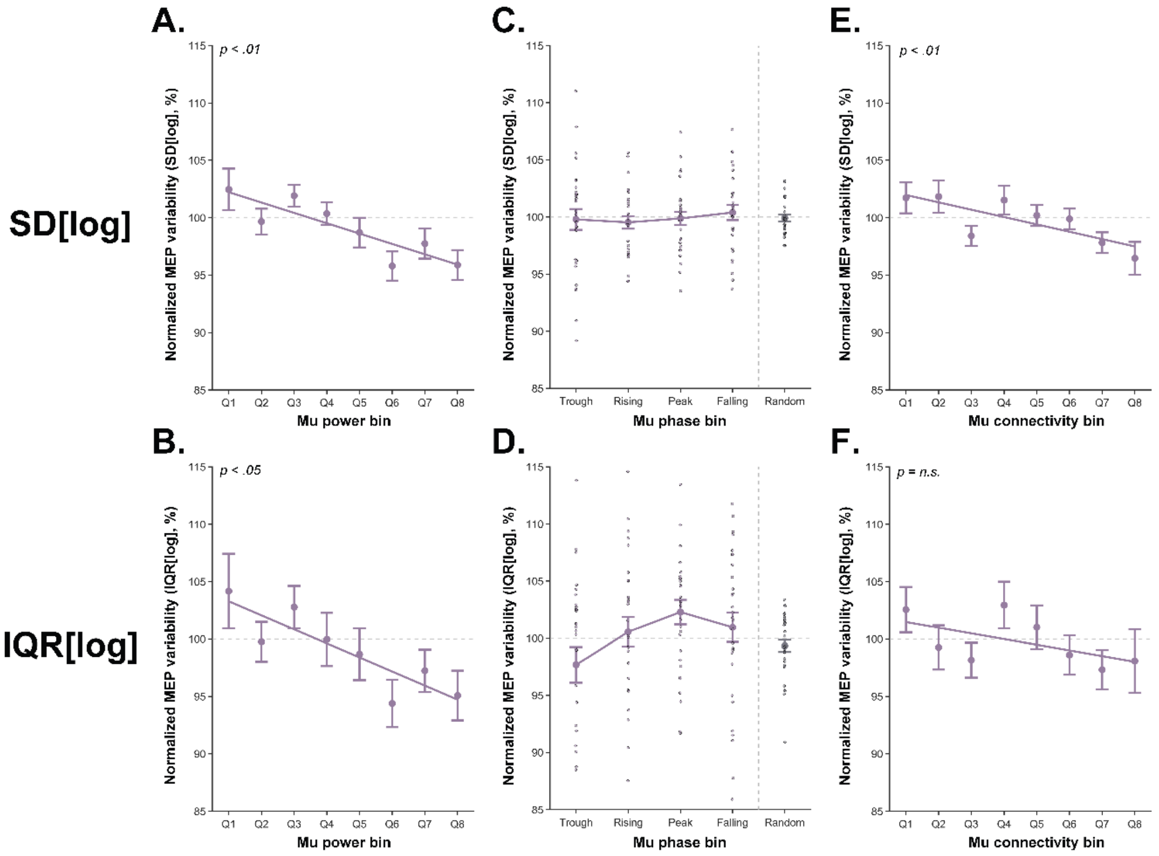


**Supplementary Figure 5. Brain-state-resolved MEP variability using alternative variability metrics.** Log transformation of MEP amplitudes reduces distributional skewness by compressing large values more strongly than small values, thereby reducing the influence of outliers and non-normal amplitude distributions. To further assess whether the observed effects were independent of mean-amplitude scaling, variability was additionally quantified using the standard deviation (SD) and interquartile range (IQR) of log-transformed MEP amplitudes [9]. These complementary variability metrics do not rely on normalization by the mean, unlike the coefficient of variation (CV). (A-B) Trial subsets with increased mu power were associated with reduced MEP variability (SD[log]: mean ± SEM slope = –0.90 ± 0.31; t(28) = –2.94, p = .007; IQR[log]: mean ± SEM slope = –1.22 ± 0.51; t(28) = –2.39, p = .024). (C-D) No significant effect of mu phase on MEP variability was observed (SD[log]: F(2.42, 67.63) = 0.20, p = .856; IQR[log]: F(2.54, 71.07) = 1.77, p = .169). (E-F) Trial subsets with increased interhemispheric M1–M1 connectivity were associated with reduced MEP variability only using SD[log] (mean ± SEM slope = –0.64 ± 0.22; t(28) = –2.85, p = .008), but not IQR[log] (mean ± SEM slope = –0.50 ± 0.40; t(28) = –1.24, p = .224). Y-axes show normalized MEP variability quantified using the SD (A-C) or IQR[log] (D-F). Variability for each state-defined bin is expressed as a percentage of total unstratified variability. The dashed horizontal line at 100% indicates the total unstratified MEP variability (SD or IQR[log] across all trials) against which within-state variability was normalized. Dots represent data from individual participants. Error bars represent ± SEM.

**Cross-site replication of brain-state-resolved MEP variability with normalized CV using validation dataset from Aalto University (Finland)**

To assess the generalizability of the findings, a validation dataset acquired at Aalto University (Finland) was included that applied a similar TMS–EEG protocol over the left M1. The study was approved by the local ethical committees of Helsinki University Hospital: HUS/1198/2016). Both research groups (Hertie Institute for Clinical Brain Research, University of Tübingen, Germany; Department of Neuroscience and Biomedical Engineering, Aalto University, Finland) are part of the ConnectToBrain Synergy Project (European Research Council, grant number 810377).

**Methods**

The validation dataset comprised 22 healthy, right-handed participants (10 female, 12 male, mean age = 27.7 ± 7.0 years). For the Aalto dataset, single biphasic pulses were delivered using a Nexstim NBS 5.2.4/NBT 2.2.4 system (Nexstim Plc, Finland). Individual T1-weighted magnetic resonance images (MRI) were used for neuronavigation with a Nexstim system (Nexstim Plc, Finland). The TMS-compatible EEG cap (EasyCap, Germany) consisted of 64 (Finland) channels and Ag/AgCl ring electrodes. Channels were online referenced to mastoid and the ground was positioned on the zygomatic bone. The randomized inter-trial interval (ITI) was 4.0–4.5 s. For further details on site-specific differences, please refer to Ahola et al. [10]. The mean RMT was 51.9% ± 9.4%. For each participant, the muscle exhibiting the highest mean response or superior signal quality was selected for subsequent analyses (FDI = 15; APB = 7).

**Results**

A significant negative linear relationship was observed between mu power and MEP variability in the Aalto dataset (mean ± SEM slope = –1.42 ± 0.59, $t(21) = -2.40$, $p = .026$; Supplementary Figure 6A) and the pooled dataset (mean ± SEM slope = –1.53 ± 0.34, $t(50) = -4.49$, $p < .001$; Supplementary Figure 6D), indicating a progressive reduction in MEP variability in trial subsets with increased mu power.

Mu phase did not significantly modulate MEP variability in the Aalto dataset ($F(2.45, 48.95) = 0.54$, $p = .624$; Supplementary Figure 6B) nor in the pooled dataset ($F(2.81, 137.77) = 0.28$, $p = .825$; Supplementary Figure 6E).

A significant negative linear relationship was observed between interhemispheric M1–M1 connectivity and MEP variability in the pooled dataset (mean ± SEM slope = –0.69 ± 0.19, $t(50) = -3.56$, $p < .001$; Supplementary Figure 6C). In the Aalto dataset (mean ± SEM slope = –0.52 ± 0.31, $t(21) = -1.68$, $p = 0.108$; Supplementary Figure 6F) it was insignificant.

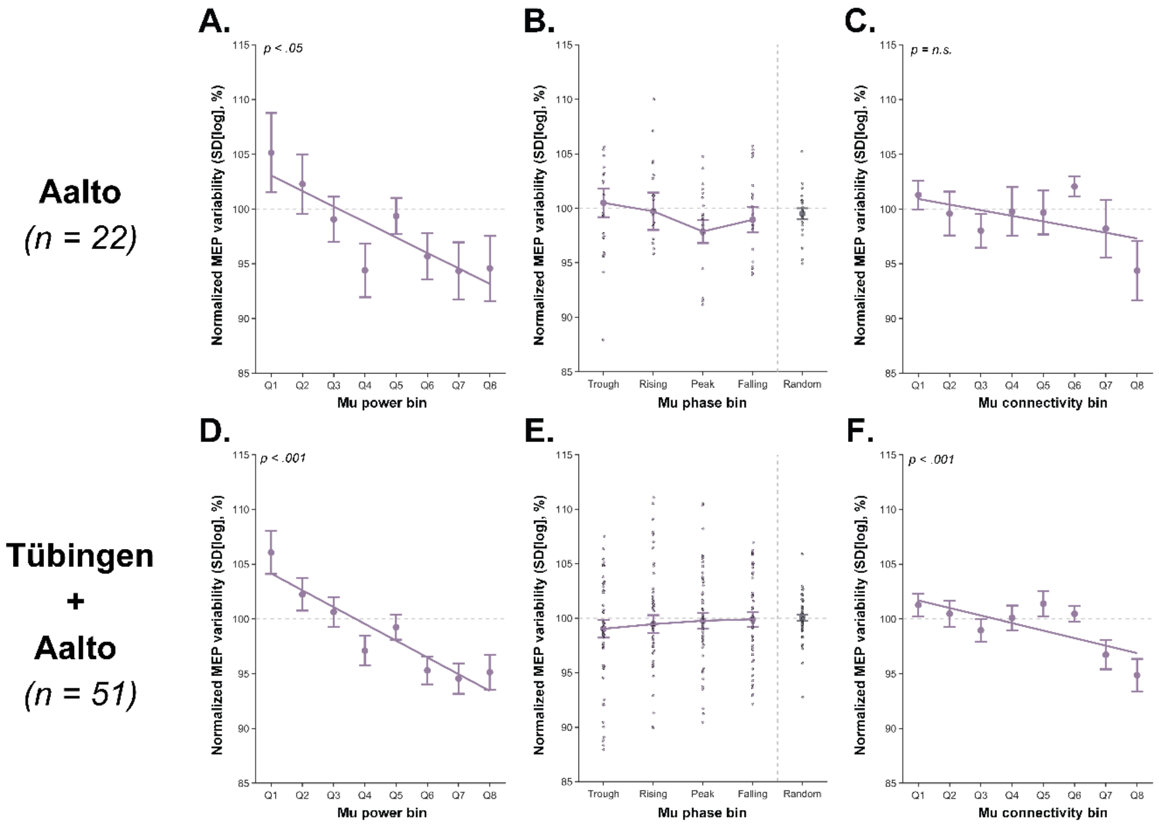


**Supplementary Figure 6. Independent replication of brain-state-resolved MEP variability (normalized CV) using validation dataset and pooled datasets. (A-C)** Results from the independent Aalto dataset showing the relationship between brain states and MEP variability across conditions. Consistent with the main Tübingen dataset, mu power stratification had a significant effect on MEP variability, with high mu power associated with reduced MEP variability. The effect of connectivity on within-state variability was not replicated. (D-F) Results from the pooled Tübingen and Aalto datasets showing the relationship between brain states and MEP variability across conditions. The effects of power and connectivity stratification on MEP variability were preserved when pooling datasets, confirming the robustness and generalizability of the observed brain-state-resolved effects. The dashed horizontal line at 100% indicates the total unstratified MEP variability (baseline CV across all trials), against which within-state variability was normalized. Dots represent data from individual participants. Error bars represent ± SEM.

**Supplementary Table 1. Descriptive statistics of effect of pre-stimulus brain states defined by mu power, phase, and connectivity on MEP amplitude and MEP variability.** Mean MEP amplitudes (µV) and mean MEP variability metrics are reported for each state and bin. Absolute variability is expressed as the standard deviation (SD[raw]). Relative variability is expressed using the coefficient of variation (CV[raw]) or standard deviation of log-transformed MEPs (SD[log]). Normalized measures are normalized to the total MEP variability and expressed as a percentage. State definitions include power (Low, High, Random), phase (Trough, Rising, Peak, Falling), connectivity (Low, High, Random), and their interactions (Power × Phase, Power × Connectivity).

| **Brain state** | **Bin** | **Mean MEP (µV)** | **Absolute variability (SD[raw])** | **Normalized absolute variability (SD[raw])** | **Relative variability (CV[raw])** | **Normalized relative variability (CV[raw]/CVtotal[raw])** | **Relative variability (SD[log])** | **Normalized relative variability(SD[log]/SDtotal[log])** |
|---|---|---|---|---|---|---|---|---|
| **Power** | Low | 1079.9 | 883.0 | 97.9 | 0.84 | 104.51 | 0.89 | 101.31 |
| | High | 1179.3 | 905.2 | 100.7 | 0.77 | 95.70 | 0.86 | 97.03 |
| | Random | 1119.2 | 896.9 | 99.4 | 0.80 | 99.88 | 0.88 | 99.93 |
| **Phase** | Trough | 1161.9 | 912.3 | 101.1 | 0.79 | 97.85 | 0.89 | 99.80 |
| | Rising | 1141.6 | 904.6 | 99.2 | 0.79 | 99.09 | 0.89 | 99.28 |
| | Peak | 1094.9 | 881.7 | 97.9 | 0.81 | 100.91 | 0.89 | 99.86 |
| | Falling | 1119.8 | 906.2 | 99.8 | 0.81 | 100.60 | 0.89 | 100.33 |
| **Connectivity** | Low | 1126.4 | 911.4 | 100.8 | 0.82 | 101.83 | 0.90 | 101.88 |
| | High | 1129.9 | 876.7 | 95.6 | 0.76 | 95.58 | 0.86 | 97.36 |
| | Random | 1133.7 | 906.6 | 99.9 | 0.80 | 99.79 | 0.88 | 99.54 |
| **Power x Phase** | Low + Trough | 1085.6 | 881.5 | 97.8 | 0.83 | 103.35 | 0.90 | 101.89 |
| | Low + Rising | 1095.3 | 903.5 | 97.9 | 0.84 | 104.55 | 0.89 | 100.85 |
| | Low + Falling | 1069.3 | 860.4 | 95.9 | 0.82 | 102.68 | 0.88 | 98.75 |
| | Low + Falling | 1065.2 | 852.9 | 95.9 | 0.84 | 104.30 | 0.88 | 101.73 |
| | High + Trough | 1240.5 | 936.9 | 102.3 | 0.76 | 92.95 | 0.85 | 94.86 |

| | | | | | | | | |
|---|---|---|---|---|---|---|---|---|
| | High + Rising | 1202.5 | 927.2 | 102.9 | 0.78 | 96.48 | 0.87 | 97.30 |
| | High + Peak | 1133.2 | 872.8 | 96.4 | 0.78 | 96.02 | 0.86 | 96.49 |
| | High + Falling | 1145.1 | 854.1 | 97.1 | 0.75 | 94.18 | 0.86 | 96.72 |
| **Power x Connectivity** | Low + Low | 1118.9 | 894.3 | 97.7 | 0.83 | 104.06 | 0.90 | 103.11 |
| | Low + High | 1049.3 | 810.2 | 88.6 | 0.79 | 98.98 | 0.88 | 97.82 |
| | High + Low | 1197.2 | 891.9 | 99.5 | 0.78 | 95.93 | 0.86 | 97.01 |
| | High + High | 1189.1 | 858.4 | 95.2 | 0.74 | 91.30 | 0.86 | 95.22 |
